# CellHeap: A scRNA-seq workflow for large-scale bioinformatics data analysis

**DOI:** 10.1101/2023.04.19.537508

**Authors:** Maria Clicia S. Castro, Vanessa S. Silva, Maiana O. C. Costa, Helena S. I. L. Silva, Maria Emilia M. T. Walter, Alba C. M. A. Melo, Kary Ocaña, Marcelo T. dos Santos, Marisa F. Nicolas, Anna Cristina C. Carvalho, Andrea Henriques-Pons, Fabrício A. B. Silva

**Affiliations:** Scientific Computing Program,Oswaldo Cruz Foundation, Av. Brasil, 4365 - Manguinhos, 21040-900, Rio de Janeiro, Brazil; Laboratory of Innovations in Therapies, Education and Bioproducts,Oswaldo Cruz Foundation, Av. Brasil, 4365 - Manguinhos, 21040-900, Rio de Janeiro, Brazil; Computational Modeling Department, National Laboratory for Scientific Computing, Av. Getúlio Vargas, 333, Petrópolis, 15651-075, Rio de Janeiro, Brazil; Department of Informatics and Computer Science, State University of Rio de Janeiro, Av. São Francisco Xavier, 524, Maracanã, 20550-900, Rio de Janeiro, Brazil; Department of Computer Science, University of Brasilia, Campus Universitário Darcy Ribeiro, Asa Norte, Brasilia, 71670-260, Federal District, Brazil

**Keywords:** single-cell RNA-seq, bioinformatics workflow, high-performance computing, COVID-19

## Abstract

**Background:** Several hundred terabytes of single-cell RNA-seq (scRNA-seq) data are available in public repositories. These data refer to various research projects, from microbial population cells to multiple tissues, involving patients with a myriad of diseases and comorbidities. An increase to several Petabytes of scRNA-seq data available in public repositories is a realistic prediction for coming years. Therefore, thoughtful analysis of these data requires large-scale computing infrastructures and software systems optimized for such platforms to generate correct and reliable biological knowledge.

**Results:** This paper presents CellHeap, a flexible, portable, and robust platform for analyzing large scRNA-seq datasets, with quality control throughout the execution steps, and deployable on platforms that support large-scale data, such as supercomputers or clouds. As a case study, we designed a workflow to study particular modulations of Fc receptors, considering mild and severe cases of COVID-19. This workflow, deployed in the Brazilian Santos Dumont supercomputer, processed dozens of Terabytes of COVID-19 scRNA-seq raw data. Our results show that most of the workflow total execution time is spent in its initial phases and that there is great potential for a parallel solution to speed up scRNA-seq data analysis significantly. Thus, this workflow includes an efficient solution to use parallel computational resources, improving total execution time. Our case study showed increased Fc receptors transcription in macrophages of patients with severe COVID-19 symptoms, especially FCGR1A, FCGR2A, and FCGR3A. Furthermore, diverse molecules associated with their signaling pathways were upregulated in severe cases, possibly associated with the prominent inflammatory response observed.

**Conclusion:** From the CellHeap platform, different workflows capable of analyzing large scRNA-seq datasets can be generated. Our case study, a workflow designed to study particular modulations of Fc receptors, considering mild and severe cases of COVID-19, deployed on the Brazilian supercomputer Santos Dumont, had a substantial reduction in total execution time when jobs are triggered simultaneously using the parallelization strategy described in this manuscript. Regarding biological results, our case study identified specific modulations comparing healthy individuals with COVID-19 patients with mild or severe symptoms, revealing an upregulation of several inflammatory pathways and an increase in the transcription of Fc receptors in severe cases.

## 1 Introduction

In recent years, single-cell technologies, such as flow cytometry, mass cytometry, and single-cell RNA sequencing (scRNA-seq), have been increasingly used to characterize cellular and molecular heterogeneity by profiling transcriptomes at the single-cell level [1]. These technologies have produced many publicly available scRNA-seq datasets. On the other hand, many sophisticated computational tools have been developed to generate knowledge of cellular processes and pathways by processing those large volumes of data. Together, these data and tools have enabled studies of cell heterogeneity and developmental dynamics across diverse biological systems, which allows a better understanding of the underlying biological processes in organisms [2, 3] and complex diseases [1, 4, 5, 6].

Specifically, scientific workflows (or pipelines) are the primary tool used to perform computational modeling based on scRNA-seq data [7], which may involve different sequencing protocols [3, 8, 9]. However, there has yet to be a generic workflow for scRNA-seq which require mastering complex computational techniques, such as highperformance computing and sophisticated analytic data science algorithms. Currently, users must develop scripts not guaranteed to cover the requirements of executing large and complex scRNA-seq experiments. Given this context, in this paper we propose a generic bio-computation service platform that focuses on high-performance computing platform deployments.

Scientific workflows represent a set of interconnected and automated tasks of an experiment, which are executed according to their input/output dependencies [10]. To create a workflow, scientists often use an *ad hoc* strategy, writing from scratch a set of scripts that connect tasks. This approach leads to different workflows, with slightly different variations of the same idea. Consequently, reproducibility, reliability, and scalability in scientific experiments may be at stake [11].

Single-cell technologies offer considerable promise in dissecting the heterogeneity of immune responses among individuals and exposing the molecular mechanisms of diseases. In particular, these technologies have been used to study coronavirus disease (COVID-19) pathogenesis [4, 12, 13, 14, 15, 16, 17, 18, 19]. COVID-19 is an ongoing global infectious disease caused by the SARS-CoV-2 virus. Single-cell technologies have bolstered the understanding of each phase of this disease and other aspects, such as differential gene expression in the single-cell multi-omics analysis of the immune response in COVID-19 [20].

It is noteworthy that most of the workflows used in the scRNA-seq studies of COVID-19 are not flexible, are built in an *ad hoc* manner, and run in a standalone desktop [14, 15, 16, 17] or on a supercomputer [8, 18].

We claim that when sophisticated bioinformatics tools are created and updated regularly, scientific workflows must be flexible, enabling users to choose between tools that execute the same task. In addition, workflows should be extensible, allowing new modules or phases to be integrated. Moreover, a workflow should also be portable, executable on a standalone platform, such as a desktop server, or on a high-performance computing platform, such as a supercomputer.

We present CellHeap, a flexible, extensible, and portable platform for scRNAseq analysis, which processes hundreds of Gigabytes of scRNA-seq raw data on supercomputer environments.

The CellHeap platform is composed of five phases. In each one, CellHeap includes many computational tools and couples them such that inputs and outputs consume/generate data in a flow that meets requirements for subsequent phases of analysis. CellHeap supports the construction of different workflows according to the goals of each research project and the associated biological question. Also, many datasets are available in public repositories, and CellHeap supports several input data formats and sequencing platforms.

We deployed CellHeap in a case study with COVID-19 data on the Santos Dumont^1^ (SDumont) supercomputer. Initially, we selected and curated publicly available COVID-19 scRNA-seq raw datasets originally processed in a sequential workflow. Then, based on the sequential workflow execution, we analysed data storage, studied computational performance, and proposed a strategy to improve execution time. In more detail, using scRNA-seq datasets of COVID-19 [4], we parallelized the workflow’s initial phases, which are the most time-consuming. Results such as the reduction in the execution time of CellHeap’s first stages for the samples representing the severe COVID-19 cases from five days (sequential) to one day and one hour (parallel) demonstrate that a simple parallel solution has great potential to speed up the workflow.

Furthermore, we examined the results produced from the workflow, designed to study particular modulations of Fc receptors in SARS-CoV-2 infection. The objective was to analyze the transcription of Fc receptors and their signaling pathways, besides cytokines/chemokines in macrophages obtained from bronchoalveolar lavage fluid (BALF) of healthy individuals and patients with mild and severe symptoms of COVID-19.

This paper is organized as follows: Section 2 introduces the CellHeap platform. Section 3 describes our case study, which deployed CellHeap in the Santos Dumont supercomputer (SDumont) considering scRNA-seq datasets of COVID-19. Section 4 presents the results obtained from the workflow using COVID-19 datasets, discusses the time and space storage needed to execute the workflow on a high-performance platform, and the results obtained are investigated from a biological perspective. Finally, Section 5 concludes this article and presents future work.

## 2 The CellHeap platform

This section describes the flexible, extensible, portable, and robust CellHeap platform for scRNA-seq customized bioinformatics workflows. CellHeap includes quality control to ensure correct results and relies on high-performance computational support for parallelizing and distributing computational tasks.

CellHeap comprises five phases (dashed rectangles in Figure 1): sample curation; data pre-processing; quality control; dimensionality reduction and clustering analysis; and advanced cell level and gene level analysis. An optional phase, sample aggregation, can also be deployed into the CellHeap workflow. The CellHeap source code is available at https://github.com/FioSysBio/CellHeapRelease.

**Fig. 1.**
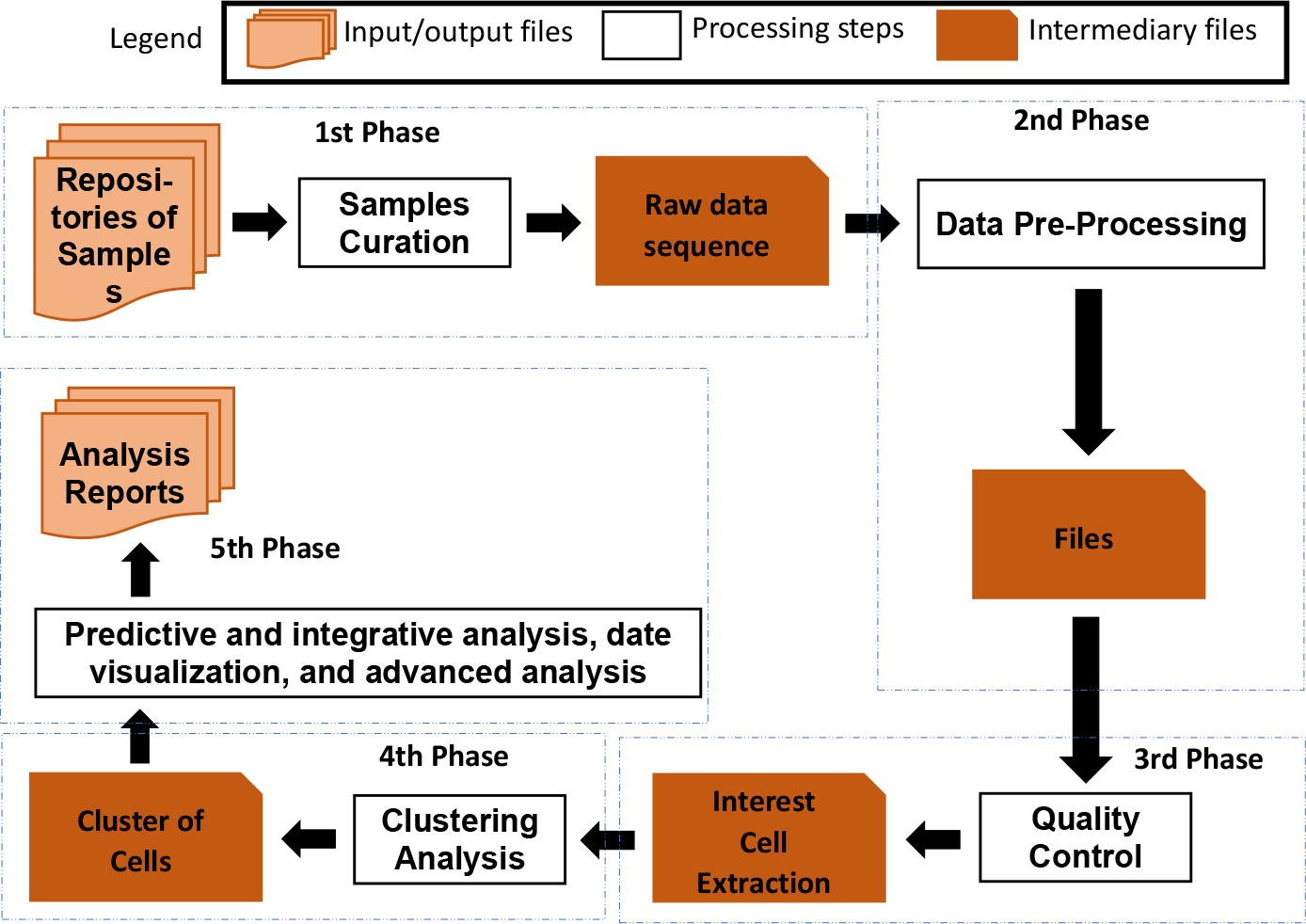
CellHeap platform conceptual view. Dashed rectangles identify the five phases of the workflow. From top-left rectangle and following the arrows, we have Phase 1: Samples Curation; Phase 2: DataPre-Processing; Phase 3: Quality Control; Phase 4: Clustering Analysis; Phase 5: Predictive and Integrative Analysis

The definition of the scientific questions under investigation is the starting point to develop experiments considering scRNA-seq datasets. The following information is essential in defining the workflow:

1. which species are considered in the research;
2. what is the origin of the tissues samples;
3. are there sufficient datasets to support the research?;
4. are the datasets available in open data repositories to guarantee reliability and reproducibility?

### 2.1 The CellHeap phases

The CellHeap platform is fully configurable to the scientific question under investigation. In this section, we detail the phases of the CellHeap platform, along with softwares that have been widely used in recent years. Chosen software in each phase of CellHeap can always be replaced if another choice is more suitable for a specific situation.

#### 2.1.1 Phase 1 - Sample curation

Public repositories, such as the Gene Expression Omnibus (GEO), contain specific samples from a single cell (e.g., tissue sample). Applying a pre-filter protocol to the scRNA-seq samples, based on several inclusion/exclusion criteria, indicates their suitability. For example, the protocol of the Expression Atlas of the European Molecular Biology Laboratory (EMBL)^2^ considers criteria containing descriptive information about a sample, such as: links to supplementary files and the sample’s raw data; verification on the series samples’ species; description of the experiment, protocol, cell line information, cell type, and disease listed by Experimental Factor Ontology (EFO) or publication data; Verification on the protocols used in the scRNA-Seq experiments (Smart-seq2, Smart-like, Drop-seq, Seq-well, 10xV2 (3 and 5 prime), and 10xV3 (3 prime)).

Since this work only includes samples that fulfill all criteria, the SRA toolkit has tools to download and validate datasets. The prefetch tool downloads the selected raw sequence data in “*.sra*” format, and samples can be identified by the “*SRR* prefix + ID from GEO”. The vdb-validate tool authenticates the successfully downloaded data. Finally, samples are converted into “*.fastq*” format files using the fasterq-dump tool according to the required input in the next phase.

#### 2.1.2 Phase 2 - Data pre-processing

This phase creates a gene count matrix from the raw sequencing data. Cellranger^3^, a package of 10x Genomics Chromium [21], is widely used for read mapping, cell demultiplexing, and to generate the cell-wise unique molecular identifier (UMI)-count table.

The cellranger count software is designed to align reads with a reference genome. Cellranger count receives the compressed “*.fastq*” input files and a reference genome to output the barcode matrix feature. There are many alternatives to cellranger count software that can replace it in CellHeap platform, for instance, UMI-tools^4^ [22] and zUMIs^5^ [23].

#### 2.1.3 Phase 3 - Quality control

The quality control (QC) phase removes artifacts to ensure that the processed cells are single and not experimentally damaged. It is a fundamental procedure in order for cell data to be used in the following phases.

Quality control for filtering sequenced cells can be performed using Seurat and SoupX packages. Seurat [24] is an R package designed for QC analysis and exploration of single-cell RNA-seq data. Seurat aims to enable users to identify and interpret sources of heterogeneity from single-cell transcriptomic measurements and to integrate diverse types of single-cell data. In Phase 3, we perform QC analysis. SoupX [25] estimates a fraction of contamination (*rho*) in the samples (reads) through the “*autoEstCont* ” function. The parameter *rho* varies from 0 to 1, where 0 indicates no contamination and one indicates 100% contamination of UMIs in a droplet. The input matrix can be corrected based on the *rho* estimation using the “*adjustCounts*” function.

#### 2.1.4 Phase 4 - Clustering analysis

This phase contains the following steps:

- Data integration;
- Dimensionality reduction of scRNA-Seq data related to cellular expression profiles [26, 27];
- Clustering of cell-level in scRNA-Seq;
- Cell-type annotation by identifying cell groups based on gene expression profiles.

Seurat’s Canonical Correlation Analysis (CCA) [28] is a tool designed for data integration. It is recommended in a recent benchmark evaluation [29] due to the algorithm’s good performance results on the batch-effect correction.

CellHeap combines UMAP [30] and PCA [31] algorithms for dimensionality reduction and cluster visualization. UMAP is a nonlinear reduction method that balances the capture of local similarity and the preservation of global data structure. Theoretically, MAP saves more time and computational cost than other algorithms, such as t-SNE, which has been the default method for such tasks in past years. However, UMAP does not preserve the cell-to-cell similarity distance, which justifies using PCA with several dozen principal components for dimensionality reduction, particularly for tasks like pseudo-time inference.

For clustering of cell-level in scRNA-Seq, one option is Metacell [32]. Metacell clusters cells with homogeneous transcriptional profiles into groups (also called metacells), and those cells that belong to a metacell have likely been resampled from the same cell. Metacell requires as input data the feature gene set. Feature gene sets may be defined by users or automatically calculated based on the gene markers with high variance. Another clustering method is provided by Seurat [33], which assigns cells to clusters based on their PCA scores. PCA scores are computed based on the expression of the most variable and previously integrated genes, generating principal components that represent a “metagene” created by combining information across a correlated set of genes.

Tools currently available for cell-type annotation include Cellassign, ScType, SingleR, and scCATCH. Cellassign [34] identifies the cell type based on a predefined set of cell markers. It implements the expectation-optimization algorithm for cell type identification using the Adam stochastic optimizer [35], executed using the TensorFlow package [36]. Cellassign can be coupled with ScType [37] in order to compensate for the potential lack in support for negative markers. SingleR [38] performs cell type annotation based on reference transcriptomic datasets of single-cell RNA from pure cell types. SingleR independently infers the cell of origin of every single cell on the target dataset. The single-cell Cluster-based Annotation Toolkit for Cellular Heterogeneity (scCATCH) [39] uses cluster marker genes in cell type annotation by matching marker genes on a taxonomy reference database of a tissue-specific cell. CellMatch is a scCATCH function that includes a panel of 353 cell types and related 686 subtypes associated with 184 tissue types and 2,097 references from humans and mice.

#### 2.1.5 Phase 5 - Advanced cell-level and gene-level analysis

The cell-level analysis focuses on the identification, characterization, and dynamics inference of groups of cells. The gene-level analysis focuses on the analysis of gene set expression among groups of cells. The choice of tools in this phase depends on the scientific question and data type under investigation.

CellHeap includes two methods for cell-level analysis: trajectory analysis, and cell-cell communication:

1. trajectory analysis: infers gene expression dynamics to capture transitions between cell identities and branch differentiation using pseudotime-based algorithms. It computes paths through cellular space that minimizes transcriptional changes among neighboring cells. A pseudotime variable describes the evolution of transcriptional states along the trajectory under certain conditions, for instance, developmental time [40, 41, 42].
2. cell-cell communication (CCC): several tools are available to infer CCC using scRNA-seq data [43]. Communication between cells usually depends on ligand-receptor interactions. First, we quantify Ligand-Receptor Interactions (LRI) coexpression; then various LRI databases can be queried, such as CellPhoneDB [44], CellTalkDB [45], or ICELLNET [46]. Other CCC methods are also associated with the fast growth of spatial transcriptomics, integrating scRNA-seq data with spatial transcriptomics or image data. They are CellTalker [47], CellChat [48], and Squidpy [49].

The CellHeap platform allows implementation of several methods for gene-level analysis, such as differentially expressed genes (DEG), gene set enrichment, pathway analysis, and metabolic analysis, as follows:

1. DEG analysis: in scRNA-Seq, gene expression variations between two conditions can be studied. The expression variation is counted in clusters of single cells, enabling marker gene identification or cell type annotation. As a general recommendation, differential expression requires non-corrected input matrices for batch effects or input matrices that incorporate technical covariates [26].
2. gene set enrichment analysis: DEGs can be grouped based on involvement in common biological processes. Some possible tools are Gene Ontology (GO), Kyoto Encyclopedia of Genes and Genomes (KEGG), the Molecular Signatures Database (MSigDB) [50], and the Reactome repository [51].
3. pathway analysis (networks of interacting genes): identifies and visualizes the main pathways involved in biological processes. CellHeap uses available resources, such as the PANTHER database [52], the Database for Annotation, Visualization and Integrated Discovery (DAVID) [53], the Reactome pathway analysis [51], and ReactomeGSA [54]. Tools for Regulon inference and TF activity prediction can also be deployed in this analysis.
4. Metabolic analysis: tools can be either pathway-based or Flux Balance Analysis-based (FBA), a standard technique for analysing genome-scale metabolic networks. Examples of FBA-Based tools are single-cell Flux Balance Analysis (scFBA) [55], Compass [56], and scFEA [57]. On the other hand, scMetabolism [58] is a pathway-based analysis tool that supports the quantification and visualization of metabolism at the single-cell resolution using scRNA-seq data.

## 3 Case study of CellHeap: COVID-19 scRNA-seq workflow

High performance computational platforms such as supercomputers are adequate for single-cell RNA-seq data analyses which require a lot of storage and processing capacity.

This section describes a case study in which the CellHeap was deployed in the Santos Dumont (SDumont) supercomputer environment using COVID-19 scRNA-seq datasets. This study aimed to analyse the modulations of Fc receptors in lung lymphoid and myeloid cells imposed by COVID-19 by processing bronchoalveolar lavage fluid (BALF) samples. We found particular modulations considering mild and severe cases.

We focus on deploying the first four phases of the workflow, which are enough to justify using high-performance environments and obtain relevant biological results. We observe that the first two phases require longer execution time and greater memory capacity when compared to the other phases.

First, we describe a sequential version of CellHeap, observing the amounts of memory, storage, and execution time for each phase of this case study. Then, we propose the deployment of a parallel version of the CellHeap platform that aims to reduce total execution time, to be detailed later.

### 3.1 The Santos Dumont computational environment

The Santos Dumont supercomputer (SDumont^6^) has processing capacity of 5.1 Petaflop/s, and presents a hybrid configuration of computational nodes. It has 36,472 CPU cores distributed across 1,134 computing nodes, the majority of which are composed exclusively of multicore CPUs. There are two categories of computing nodes, base and sequana. The base nodes have 64GB RAM and are connected using an Infiniband FDR (56Gb/s) network. The sequana nodes have 384 GB or 768 GB RAM and are connected using an Infiniband EDR (100Gb/s) network. There are 94 GPU sequana nodes with NVIDIA Volta V100 accelerators and 384 GB RAM. Additionally, SDumont has the Lustre parallel file system with a capacity of 1.7 PBytes integrated with the Infiniband network, and a secondary file system of 640 TBytes.

The SDumont Supercomputer uses the Slurm^7^ as its cluster workload manager. The Slurm is an open-source cluster management and job scheduling system for Linux clusters. The SDumont has 26 queues to submit jobs with different priorities to attend to workload characteristics.

The size of the raw sequence data of each scRNA-seq sample varies from dozens to hundreds of gigabytes. Therefore, choosing one of the execution queues depends on the raw sequence data size. Our study used the SDumont queues composed of sequana nodes with 384 GB RAM (sequana cpu queue), or 768 GB RAM (sequana cpu bigmem queue).

### 3.2 Sequential Execution of CellHeap

This section lists the main tools used in our case study, created on the CellHeap platform and deployed on the SDumont supercomputer. It uses the SRA toolkit in the first phase and R packages in the following phases.

First off, at SDumont, the large and variable-sized input samples led to the need for an evaluation of memory consumption and execution time in all workflow phases. If memory capacity or execution time is misestimated, the SDumont Slurm scheduler will not execute the task to completion, killing it upon attaining the maximum execution time for the assigned queue.

We began with sequential execution of the workflow in order to estimate memory consumption and execution time, taking into consideration the multithreading capacity of some tools. This way, we determined which SDumont submission queues meet requirements for both factors to process the input samples fully.

#### 3.2.1 Phase 1 - Sample curation and download

In this case study, CellHeap accessed the GEO [59] repository to collect samples that meet criteria defined by the EMBL^8^ protocol. From the SRA toolkit^9^, CellHeap used four tools to manipulate samples: i) vdb-dump to obtain information; ii) prefetch to download the SRA samples; iii) vdb-validate to verify if the downloaded samples are consistent; iv) fasterq-dump to convert *sra* files into *fastq* files.

The workflow received as input the bronchoalveolar scRNA-seq dataset PRJNA608742 from the NIH GEO repository^10^. This dataset was generated by Liao et al. [4] and considered the samples of 12 patients as input data. Three were collected from healthy people (the control samples), three samples were collected from patients with mild COVID-19 symptoms, and six were from patients with severe COVID-19 symptoms. However, after curation and quality control, we processed only 11 samples. We downloaded raw data samples from NCBI SRA. The vdb-dump, prefetch, and vdb-validate tools have sequential implementations, and do not offer the possibility of parallelization.

The fasterq-dump tool produced the input files for Phase 2. It is a multithreaded tool that consumes significant temporary disk space in addition to the final output file. We noted that at least 13 times the sample size was needed in disk storage to guarantee the correct completion of this tool execution.

Each sample had two read files for paired-end runs. Due to variable input file sizes, Phase 1 processing used the cpu big mem queue, which has computing nodes with 768 GB RAM.

Two scripts were submitted to run this phase: one to download the input data (prefetch tool) sequentially, and the other (multithreaded) to generate the fastq input files for Phase 2 (fasterq-dump tool).

Phase 1 was the most time-consuming step of this case study. The prefetch tool downloaded the 11 raw data samples (913.1 GB) in approximately 16 hours. However, after some tests, we realized that the limit imposed on the network bandwidth came from the public repository itself. The following tool executed in Phase 1 is fasterq-dump. It is a multithreaded tool that took 155 hours for the 11 samples. Therefore, the total execution time was approximately 172 hours for Phase 1, considering 11 samples. After evaluating execution times and tools, we noted that fasterq-dump execution might be an opportunity for parallelization.

#### 3.2.2 Phase 2 - Reads mapping and quantification

In Phase 2, Cellheap used Cellranger count (v.6.1.0) to align reads to a reference genome for each sample’s file. Cellranger count processes scRNA-Seq data and generates the gene-barcode matrix, demanding a large amount of storage and processing capacity. This task produced several output files, of which the barcodes.tsv, matrix.mtx, and features.tsv, the input files for Phase 3, are the most representative. The output datasets generated by Cellranger count also provide quality assessment reports that display essential quality metrics (i.e., Q30).

In Phase 2, the appropriate use of available computing power is crucial to efficiently execute our scRNA-Seq workflow with COVID-19 data. This phase also required submission queues that have computing nodes with 768 GB RAM.

We assembled a hybrid reference genome to align reads for this case study. The assembly of the hybrid genome is executed only once by invoking the Cellranger mkref function, which includes both the human genome (GRCh38 v3.0.0) and the SARSCoV-2 genome (NC 045512v2) of the severe acute respiratory syndrome coronavirus 2 isolate genome Wuhan-Hu-1. The reference genome was built offline and was an input parameter for Cellranger count.

Notably, Phase 2 processing was both data and computing-intensive. For example, cellranger count required more than 104 hours to process all samples in a single computational node. In the worst case, one control sample took more than 20 hours to be processed. Processing time could be improved by distributing computing loads among multiple nodes.

#### 3.2.3 Phase 3 - Quality control

In Phase 3, CellHeap used R (v.4.2.1) [60], an integrated suite of software facilities for data manipulation, calculation, and graphical display. Many packages are available through the CRAN family of Internet sites providing a wide variety of statistical and graphical routines. CRAN is a network of FTP and worldwide web servers that stores identical, up-to-date code and documentation versions for R^11^.

CellHeap performs data quality control by filtering cells. First, we used SoupX [25] v.1.6.2 to estimate ambient RNA contamination. Then, we used Seurat [24] to filter cells by removing cell barcodes according to the three criteria commonly used for scRNA-Seq quality control processing [26]: UMI counts; genes expressed per cell; and percentage of mitochondrial DNA. The filtering parameters must be refined based on the tissue and the scientific question under investigation. After filtering, we remove cells contaminated with the SARS-CoV-2 virus and use only non-contaminated cells in the downstream analysis.

Phase 3 executed quickly in comparison to previous phases, taking from 1 to 7 minutes per sample, depending on its size. For all 11 samples of the PRJNA608742 input dataset, the total sequential execution time was 26 minutes. Phase 3 required less memory than previous phases, and used the sequana nodes (384GB RAM) for submission queues at the SDumont. The output files were *tsv* files that serve as input to Phase 4. Parallelizing Phase 3 would offer only marginal gains to the CellHeap workflow so we decided to keep Phase 3 as a sequential workflow step.

#### 3.2.4 Phase 4 - Sample integration, cell clustering, and cell annotation

Phase 4 included software tools for ScRNA-seq dataset integration, dimensionality reduction, cell clustering, and cell annotation based on gene markers and gene expression profiles.

In this phase, we used the following tools: Seurat for data integration, dimensionality reduction, and cell clustering. In contrast, Cellassign (version 0.99.21) and ScType to identify the cell type based on a predefined set of gene markers.

Seurat carries out the clustering through a graph-based approach on a PCA reduction with several dozens of PCs (typically *≥* 30). Based on the Euclidean distance in the PCA space, we first constructed a KNN graph then used the Louvain algorithm to cluster the cells. Finally, we used the UMAP algorithm for cluster visualization, with the PCA reduction as input.

In Phase 4, we grouped samples depending on level of the severity. Therefore, there were three groups of integrated samples: control, mild and severe, with specific scripts for each group. Phase 4 scripts require a submission queue to nodes with 768 GB RAM. Groups can be processed concurrently.

Phase 4 had a longer execution time than Phase 3. Depending on the total number of cells per integrated condition, execution time varied from over 11 hours to less than 1 hour. The total sequential time was over 17 hours, and there is room for improvement by processing different conditions simultaneously.

### 3.3 Parallel execution of Phases 1 and 2

As stated in Sections 3.2.1 and 3.2.2, the first two phases of CellHeap are compute-intensive and take many hours to execute. In fact, those phases execute separate and independent subworkflows for each sample, as shown in Figure 2 and, as such, may be good candidates for parallelization.

**Fig. 2.**
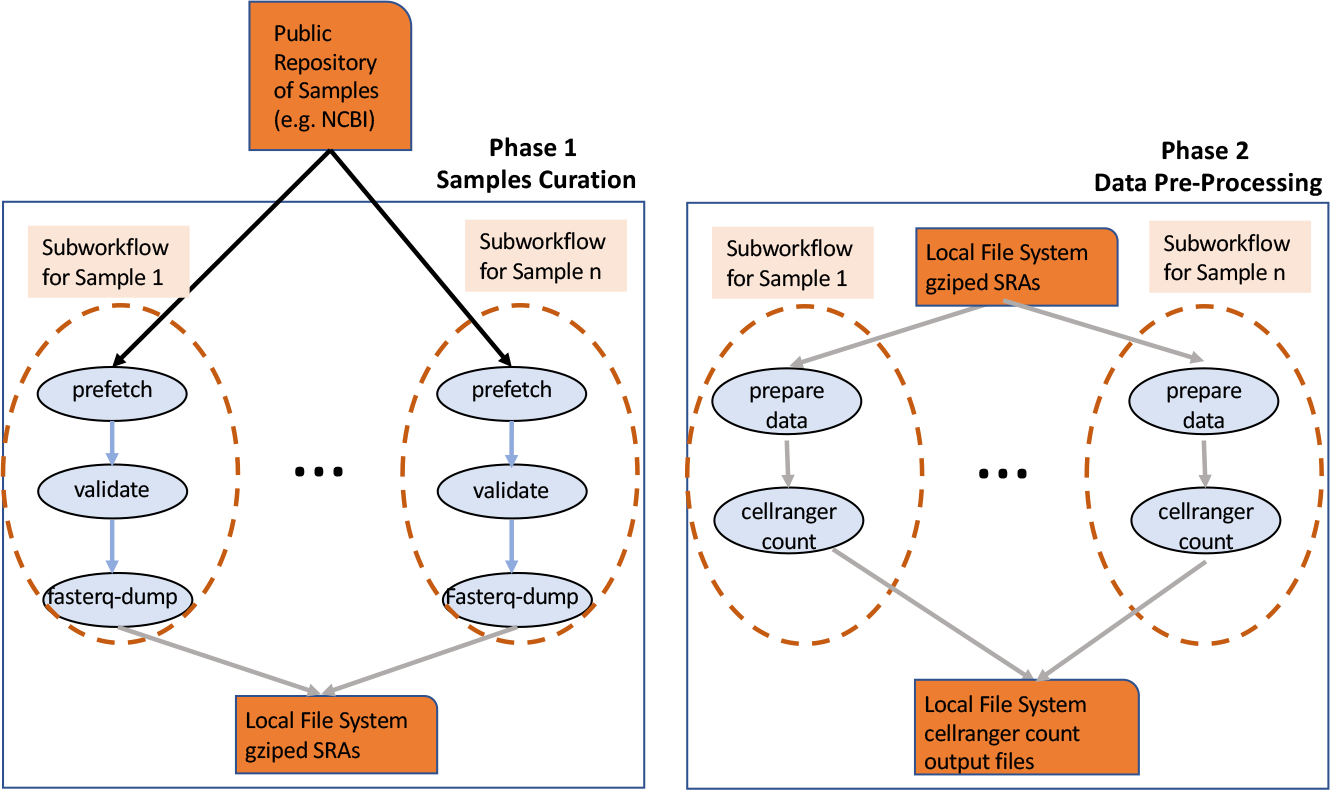
Subworkflows in Phases 1 and 2, which are candidates for parallelization.

In our design, we devised a parallel architecture that considers many computing nodes. This type of architecture is available on high-performance platforms, such as supercomputers. In this way, we can explore multinode processing to accelerate the time-consuming CellHeap phases/tools.

In Phases 1 through 3 of the sequential approach, each sample is computed by a single node, using the linear workflow shown in Figure 3, the input of one task being the output of the previous one. If, for instance, there are two samples, sample 1 will be fully processed before the processing of sample 2 begins.

**Fig. 3.**
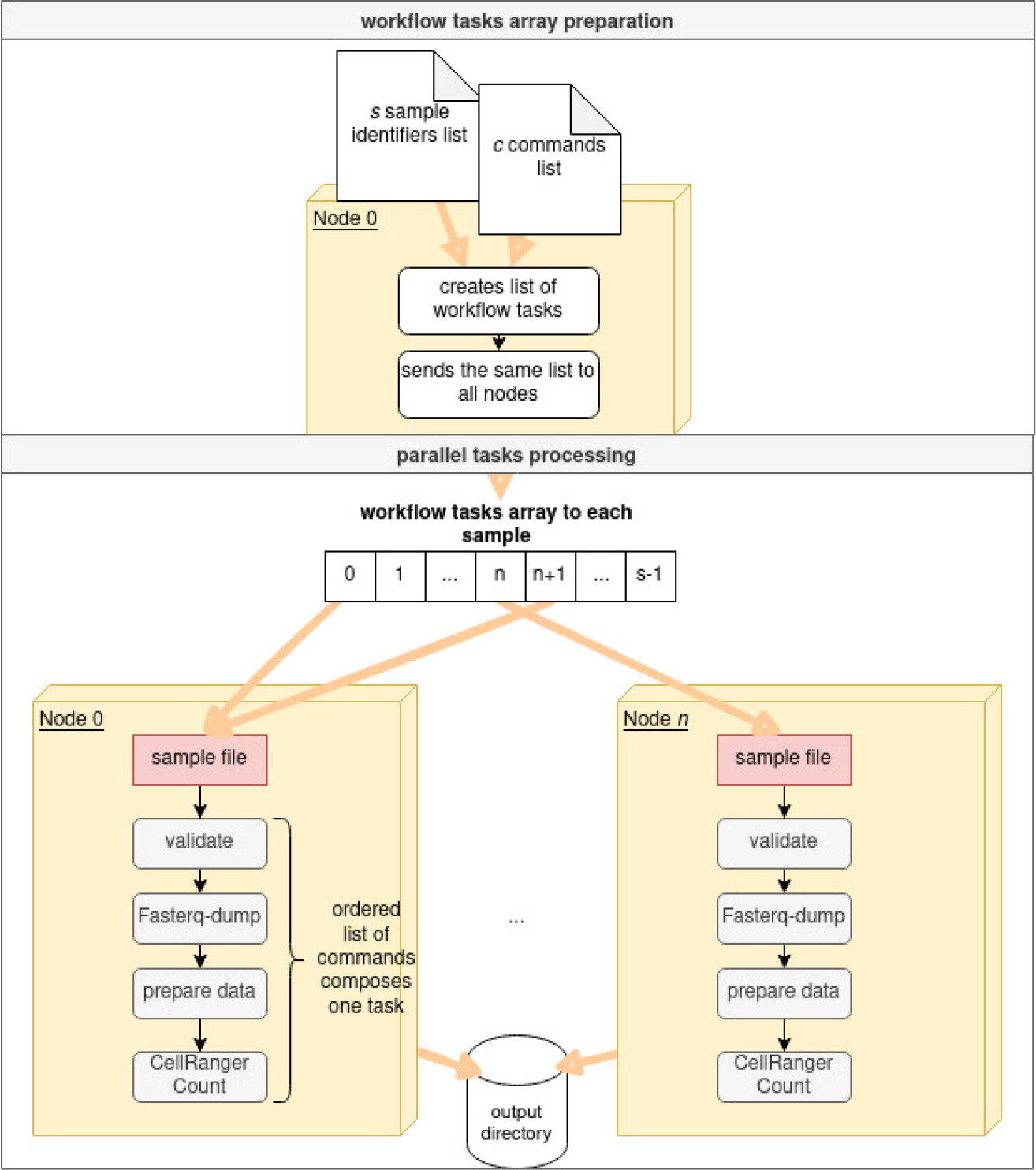
Automated parallel tool to parallelize Phases 1 and 2 of CellHeap.

However, since the processing of each sample in Phases 1 through 3 is independent of the others, the samples may be processed in parallel. We can therefore exploit more parallelism in the execution of the workflow.

Phases 1 and 2 consume the most memory storage and processing time in CellHeap.

These phases, when parallelized, can provide substantial performance gains.

The prefetch tool, executed in Phase 1, downloads samples from public repositories. Doing tests in the Santos Dumont supercomputer, we noticed that the effective bandwidth of the downloads was low. So, we contacted RNP (National Network of Education and Research), that is responsible for the Brazilian backbone and the international network connections. The RNP staff did extensive tests and discovered that the bandwidth was limited by the repository provider, in this case, NCBI. So, for the prefetch tool, parallelization will not incresase the performance significantly.

Therefore, our parallel approach focused on the other tools used in phases 1 and 2, e.g., validate, fasterq-dump, prepare data and cellranger count. Since each sample is processed independently in phases 1 and 2, i.e., there is no need to process all samples in phase 1 before starting phase 2, we designed a parallel asynchronous strategy that merges phases 1 and 2, executing the subworkflows for each sample independently, as shown in Figure 3. For instance, node *n*_0_ may be executing fasterq-dump for sample 0 and node *n*_1_ may be executing cellranger count for sample 1.

Using this list, we created a peer approach to process the samples simultaneously, as shown in Figure 3. In our automated approach, we first create *n* Message Passing Interface (MPI) processes, where *n* is the number of nodes. Since the execution time is proportional to the sample size, node 0 first sorts the samples by size and generated a list of sample ids, organized from largest to smallest. This sample ids list is sent to the *n −* 1 nodes, from 1 to *n −* 1.

Upon receiving the list, each node uses its node number to process the samples independently, avoiding synchronization. Our approach is lock-free, with no wait time for the accessing list of samples, since each node always accesses a different element of the list. Each node calculates a hash using the function *s_i_* mod *N* = *n_i_*, where *s_i_* is the sample number, *N* is the total number of nodes, *n_i_* is the node number, and mod returns the remainder when dividing *s_i_* by *N* . In Figure 3, we show *n* + 1 nodes.

Assuming that there are 2 nodes and that the number of samples *s* is equal to 8, nodes 0 and 1 will process samples (0, 2, 4, 6) and (1, 3, 5, 7), respectively. Each node will process a linear subworkflow (from validate to cellranger count).

At the end of each subworkflow computation, the nodes write the output in a shared directory. It is important to note that, in addition to being compute-intensive, fasterq-dump, gzip and cellranger count also execute a great number of I/O operations in the shared file system.

When a node finishes computing all its assigned subworkflows, it sends a message to node 0. When node 0 receives terminating messages from all nodes, the execution of phase 2 is completed.

## 4 Results and Discussion

In this section, we first discuss the obtained results related to execution time and memory. Then, we interpret the results from the point of view of their biological implications.

As we mentioned in the previous section, we executed the sequential and parallel versions of our CellHeap workflow for RNA-seq samples of project PRJNA608742, retrieved from NCBI. This dataset contains control, mild and severe cases of BALF samples.

### 4.1 Computational Results

Table 1 shows the total time obtained when executing the sequential version of the CellHeap platform, considering the 11 samples PRJNA608742. The size of all SRA files for 11 samples was 913.1 GB. The amount of storage required for processing is multiplied more than 10 times during Phase 2 execution.

**Table 1.** Execution time of the sequential version of the workflow

| Phase | Execution time<br>(hh:mm) |
| --- | --- |
| *Phase 1* | 172:13 |
| *Phase 2* | 104:18 |
| *Phase 3* | 00:26 |
| *Phase 4* | 17:15 |
| *Total time* | 294:12 |

Phases 1, 2, and 3 are executed only once. However, Phase 4 may be executed several times to adjust parameters and answer the biological question correctly. Table 1 shows the time for a single run of Phase 4.

It is important to note that a single execution of the workflow required approximately 294 hours, or 12 days, of intensive processing, considering only the 11 samples of our case study. Furthermore, the execution times do not include wait time in SDumont’s slurm scheduler.

To show the potential for parallelization in our COVID-19 CellHeap workflow, we focused on Phases 1 and 2. Moreover, we did not parallelize the execution of the prefetch tool since its performance seems to be defined by the NCBI public repository download policy. In this section, we focused on the six severe samples since they are large enough to be representative. In addition, we organized sample based on size to distribute the samples among the processing nodes efficiently.

Table 2 shows the name, size, and sequential running time of the six severe case samples for Phases 1 and 2. The tools used in these phases are validade, fasterq-dump, gzip, and cellranger count. This table shows that sequentially processing the severe samples takes about five days and requires substantial RAM.

**Table 2.** The size and sequential execution time of the six severe samples

| Sample name | Size Gb | Execution Time (hh:mm) |
| --- | --- | --- |
| SRR11537950 | 40.8 | 09:22 |
| SRR11537951 | 46.2 | 10:52 |
| SRR11537949 | 46.3 | 10:53 |
| SRR11181959 | 85.7 | 28:39 |
| SRR11181958 | 90.2 | 29:40 |
| SRR11181956 | 95.9 | 30:37 |
| Total | 405.1 | 120:03 |

To simultaneously process the samples in Phase 1 (validate, fastq-dump, gzip) and Phase 2 (cellranger count) in a balanced way considering memory consumption and running time, we first allocated three nodes for execution and, since there were six samples, each node processed two samples. The samples were sorted by their size and assigned to the nodes in a striped pattern, with a hash function. Thus, nodes 0, 1 and 2 computed samples (0, 3), (1, 4) and (2, 5), respectively. Then, we assigned six nodes for execution, each of which computed one sample. As a result, the execution times to process the severe samples (Phases 1 and 2) sequentially and in parallel (3 and 6 nodes) are shown in Table 3.

**Table 3.**
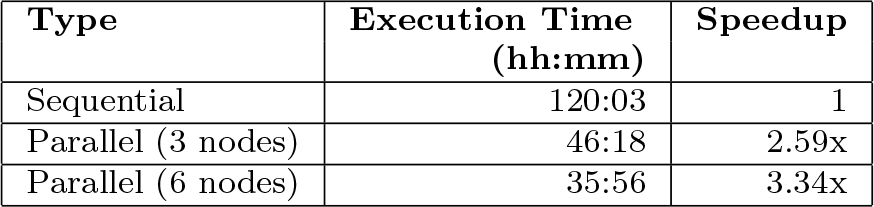
Execution time of the six severe samples (Phases 1 and 2)

| Type | Execution Time (hh:mm) | Speedup |
| --- | --- | --- |
| Sequential | 120:03 | 1 |
| Parallel (3 nodes) | 46:18 | 2.59x |
| Parallel (6 nodes) | 35:56 | 3.34x |

There is a substantial reduction in total execution time when jobs are triggered simultaneously in the parallel version. For example, processing the six samples with three nodes required one day, 22 hours, and 18 minutes. On the other hand, sequential processing of the same samples took five days and 3 minutes, achieving a speedup of 2.59x, an impressive reduction of more than three days in processing time. When we used six nodes, the execution time was one day, 11 hours, and 16 minutes, reducing execution time by 11 hours as compared to execution with three nodes. These results demonstrate the great advantage of using our parallel tool in specific phase of CellHeap.

Phase 4 poses another opportunity for parallelization, with the control, severe, and mild case sample groups being executed several times depending on the biological aspect under analysis. In this case, we executed each group (severe, mild, control) as separate jobs, as shown in Table 4. In this table, in addition to the group, we show the execution time and the number of cells processed. Note that the running time is proportional to the number of cells.

**Table 4.** Execution time of the sequential version of the workflow by group (Phase 4)

| Job | Group<br>(number of<br>samples) | Execution<br>Time<br>(hh:mm) | Number<br>of cells |
| --- | --- | --- | --- |
| 1 | Severe (6) | 11:19 | 40,733 |
| 2 | Mild (2) | 00:41 | 2,486 |
| 3 | Control (3) | 05:15 | 18,585 |
|  | Total Exec Time (sequential) | 17:15 |  |
|  | Total Exec Time (parallel) | 11:19 |  |

In Phase 4, we observe a reduced execution time when jobs are run in parallel. In this case, we reduced the execution time from 17:15 to 11:19 hours, achieving a speedup of 1.52x (Table 4).

Each group’s time presented in Table 4 corresponds to a single execution. However, in many cases, more than one run is needed, using different parameters, and each one of those runs benefit from parallelism.

### 4.2 Biological results

We analysed transcriptional modulations of Fc receptors (FcR) induced by the SARSCoV-2 infection to test and validate our tool. FcRs bind to the Fc portion of immunoglobulins (antibodies) (Fig. 4A), with some members binding to IgG (*FcγR*), IgA (*FcαR*), or IgE (*FcεR*) [61]. The engagement of the Fc domain of IgG (Fig. 4A) with members of the *FcγR* family mediates most cellular functions induced by this antibody class, which is particularly important because many IgG-based alternative therapies to treat COVID-19 are under investigation. These include the injection of neutralizing monoclonal antibodies against multiple inflammatory mediators and the passive transfer of polyclonal plasma from convalescent patients [62].

**Fig. 4.**
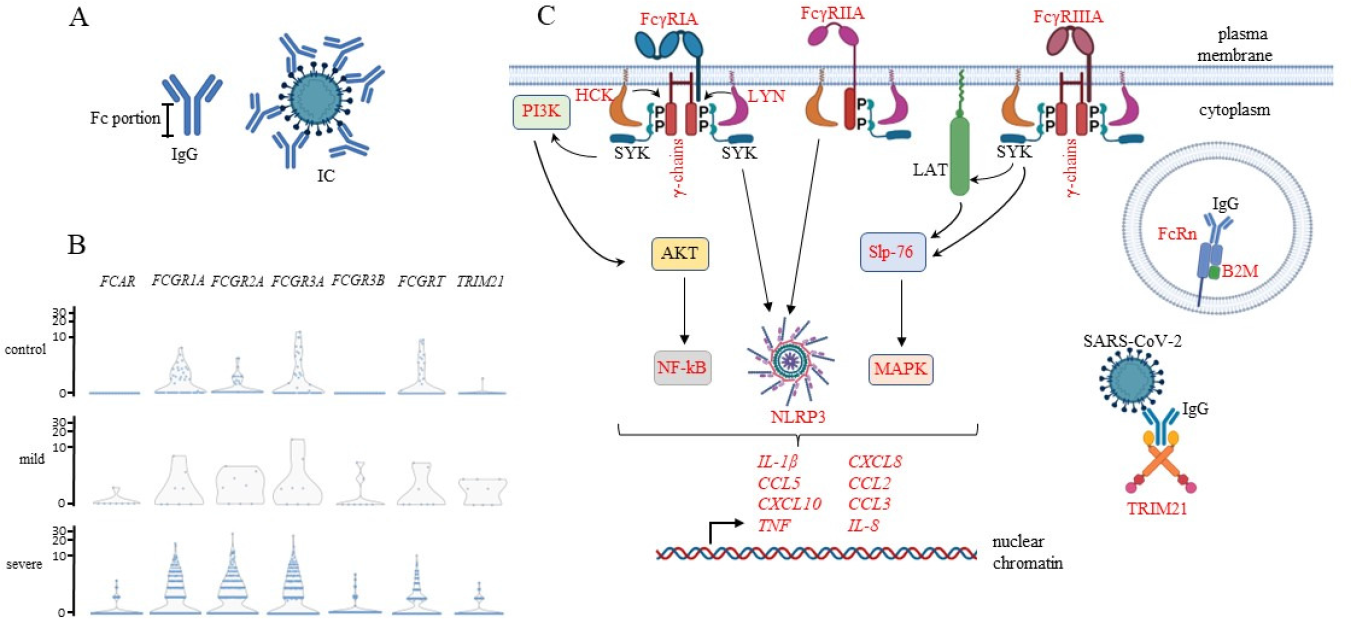
A*n*alysis *of Fc receptors in macrophages from COVID-19 patients.* The biological analysis was performed in bronchoalveolar lavage fluid (BALF) macrophages obtained from healthy (control) or COVID-19 patients with mild or severe symptoms. A) shows a schematic representation of an IgG molecule with its Fc portion and an immunocomplex (IC) composed of IgGs bound to a SARS-CoV-2 virus. B) violin plots of the indicated FcRs for the three groups of patients. C) alignment of analyzed molecules that participate in FcRs’ response according to known intracellular pathways. All molecules represented in red were upregulated in severe COVID-19 cases compared to the other groups. All three kinases, HCK, LYN, and SYK, can associate with each FcR represented and trigger the downstream signaling pathways. The pathways were separately represented only for didactic purposes.

Considering the *FcγR* family, only the *FcγR*1*A* binds with a high affinity to monomeric IgG, while *FcγR*2*A*, *FcγR*3*A*, and *FcγR*3*B* bind to multimeric IgG immune complexes (IC) or opsonized particles [63] (Fig. 4A). These interactions lead to stimulatory signals, such as phagocytosis, antibody-dependent cytotoxicity, degranulation of inflammatory mediators, cytokines release, and much more. The *FcγR*2*B* also binds to the Fc portion of IgGs composing IC, but it releases inhibitory signals when co-aggregated with other membrane receptors [64, 65]. All receptors cited are expressed on the cell membrane. TRIM21 and FcRn participate in antigen processing and, therefore, in T lymphocytes’ activation, favor anti-virus immunity. The TRIM21 binds to IgG-coated viruses (or other pathogens/molecules), leading to virus degradation by proteasomes and reduced viral load [66]. The FcRn protects IgGs in the intracellular environment, increasing their half-life and bio-disponibility [67], making it a potentially important player after IgG-based therapies. TRIM21 and FcRn participate in antigen processing for peptide presentation in major histocompatibility complexes (MHC) and T lymphocyte-based immune response, favoring anti-viral immunity.

We analysed macrophages in BALF obtained from healthy individuals and COVID19 patients with mild or severe clinical manifestations. First, the macrophages were identified based on the simultaneous transcription of the CD14, CD68, STAT1, TNF, IL6, IRF5, and IL1B genes. Then, we analyzed the transcription of FcRs, intracellular molecules involved in FcR signaling pathways, and cytokines/chemokines. The violin plots of FCAR, FCGR1A, FCGR2A, FCGR3A, FCGR3B, FCGRT, and TRIM21 transcription (Fig. 4B) showed some modulations when comparing COVID-19 severe cases with healthy individuals or those with mild symptoms. In severe cases, more macrophages transcribed FcRs (increased cellular frequency), and there was an upregulation at the transcriptional level, especially of FCGR1A, FCGR2A, and FCGR3A (upregulation per cell) (Fig. 4B).

When we arranged our findings in a coherent cellular response (Fig. 4C), we observed a scenario compatible with a prominent inflammatory response in severe cases. In Fig. 4C, we represented the molecules upregulated in severe cases in comparison to macrophages from the other groups (in red).

After ligand engagement, members of the SRC family kinases are activated [68], and we observed the upregulation of LYN and HCK in our analysis. These kinases then phosphorylate ITAM (tyrosine-based activation motifs) sequences (YxxI/L) in the intracellular tail of *FcγR*-associated *γ* chains (*FcγRIA* and *FcγRIIIA*) or in the *FcγRIIA* itself (Fig. 4C) [69]. This phosphorylation event leads to the recruitment and activation of SYK [70], which was transcribed but not upregulated in our analysis (Fig. 4C). SYK phosphorylates many target proteins, including the PI3K, which is then recruited to the membrane [71]. SYK also directly phosphorylates Slp-76 (LCP2) or phosphorylates LAT, that in turn phosphorylates Slp-76 [72, 73]. Moreover, SYK [74] and HCK [75] are essential in assembling NLRP3 inflammasome, a pivotal inflammatory structure [76]. These initial membrane-associated events trigger multiple inflammatory pathways whose components were upregulated in our analysis. The PI3K activates the AKT pathway, one of the NF-kB activators, a critical player for the transcription of numerous inflammatory mediators, such as cytokines and chemokines, that lead to the activation, differentiation, and recruitment of multiple inflammatory cell types [77]. The activation of Slp-76 leads to the activation of downstream inflammatory pathways, but we observed the upregulation of only MAPK-dependent pathways [78]. The predicted activation of these pathways is compatible with the upregulation of the inflammatory mediators we observed: TNF, IL-1*β*, CCL5, CXCL10, CXCL8, CCL2, IL-8 (Fig. 4C), and others. Finally, we observed an increase in the transcription of FcRn and its required accessory molecule B2M and TRIM 21, with yet unknown outcomes to the infection. We plan to analyse more FcR family members in other myeloid and extend to lymphoid cells to better understand the FcR’s impact on the SARS-CoV-2 infection.

## 5 Conclusion

This paper presents CellHeap, a flexible, extensible, and portable platform for scRNAseq data processing deployable in supercomputers. Bioinformatics tools for scRNA-seq are a very active area of research today, and the number of new tools has increased substantially in recent years. Furthermore, the amount of scRNA-seq data available for analysis in public repositories is growing rapidly. Therefore, flexible, robust, and scalable workflows, like the ones produced by CellHeap, are paramount to processing this massive amount of data and increasing biological knowledge.

As a case study, we designed a workflow to investigate particular modulations of Fc receptors, considering mild and severe cases of COVID-19. This workflow was deployed in the Brazilian Santos Dumont supercomputer and could process dozens of Terabytes of COVID-19 scRNA-seq raw data. Our results showed that most of the workflow total execution time is spent in its initial phases. Thus, we developed an efficient solution to use parallel computational resources, improving the total execution time of the workflow. Regarding the biological results, we emphasize the higher transcription of Fc receptors in macrophages from severe cases, besides a prominent inflammatory response associated with the upregulation of Fc receptors.

We expect CellHeap to improve continuously, aggregating new tools as they are validated and available. We also expect to soon deploy CellHeap in other high-throughput platforms, such as computing clouds.

## 6 Competing interests

No competing interest declared.

## 7 Author contributions statement

V. S. S., M. O. C. C., M. C. S. C., H. S. S., and M. T. S. implemented the workflow, V. S. S., M. O. C. C., M. C. S. C., H. S. S., F. A. B. S. conceived and conducted the experiments, K. A. C. O., M. F. N., A. C. C. C., and A. H.-P. analyzed the results, M. E. M. T. W., A. C. M. A. M., and F. A. B. S. wrote the manuscript. All the authors reviewed the manuscript.

## Acknowledgments

The authors acknowledge the National Laboratory for Scientific Computing (LNCC/MCTI, Brazil) for providing SDumont supercomputer HPC resources, contributing to the research results reported in this paper. URL: http://sdumont.lncc.br. The authors also acknowledge the INOVA-FIOCRUZ program (grant number VPPCB-005-FIO-20-2-34-52) for funding this research. M.E.M.T. Walter and M.F.N thank CNPq for the research scholarship PQ 306947/2021-8 and PQ 306894/2019-0, respectively.

## Footnotes

1. https://sdumont.lncc.br/

2. https://www.ebi.ac.uk/gxa/sc/help.html

3. https://support.10xgenomics.com/single-cell-gene-expression/software/pipelines/latest/what-is-cell-ranger

4. https://github.com/CGATOxford/UMI-tools

5. https://github.com/sdparekh/zUMIs

6. SDumont details in https://sdumont.lncc.br.

7. Slurm details in https://slurm.schedmd.com.

8. https://www.ebi.ac.uk/gxa/sc/help.html

9. https://github.com/ncbi/sra-tools

10. https://www.ncbi.nlm.nih.gov/Traces/study/?acc=PRJNA608742

11. https://cran.r-project.org/

## Notes

https://github.com/FioSysBio/CellHeapRelease

### Competing Interest Statement

The authors have declared no competing interest.

### Summary of Updates

Abstract, Results, and Conclusion

## References

[1] Tian Y, Carpp LN, and Miller HER et al. Single-cell immunology of SARS-CoV-2 infection. Nature Biotechnology, 40:30–41, 2022.

[2] Kuchina A, Brettner LM, and Paleologu L et al. Microbial single-cell rna sequencing by split-pool barcoding. Science, 2020.

[3] Vigneron A, O’Neill MB, and Weiss BL et al. Single-cell rna sequencing of trypanosoma brucei from tsetse salivary glands unveils metacyclogenesis and identifies potential transmission blocking antigens. Proceedings of the National Academy of Sciences, 117(5):2613–2621, 2020.

[4] Liao M, Liu Y, and Yuan J et al. Single-cell landscape of bronchoalveolar immune cells in patients with covid-19. Nature medicine, 26(6):842–844, 2020.

[5] Carangelo G, Magi A, and Semeraro R. From multitude to singularity: Na up-todate overview of scRNA-seq data generation and Analysis. Frontiers in Genetics, 13:994069, 2022.

[6] Zhang B, Moorlag SJ, and Dominguez-Andres J et al. Single-cell RNA sequencing reveals induction of distinct trained-immunity programs in human monocytes. The Journal of Clinical Investigation, 132(7):e147719, 2022.

[7] Hong R, Koga Y, and Bandyadka S et al. Comprehensive generation, visualization, and reporting of quality control metrics for single-cell RNA sequencing data. Nature Communications, 13(1688):1–9, 2022.

[8] Van de Sande B, Flerin C, and Davie K et al. A scalable SCENIC workflow for single-cell gene regulatory network analysis. Nature Protocols, 15:2247–2276, 2020.

[9] Shomroni O, Sitte M, and Schmidt J et al. A novel single-cell RNA-sequencing approach and its applicability connecting genotype to phenotype in ageing disease. Scientific Reports, 12(4091):1–14, 2022.

[10] Aalst WMP. Flexible Workflow Management Systems: An Approach Based on Generic Process Models. In Proceedings of the Database and Expert Systems Applications (DEXA), pages 186–195, 1999.

[11] Deelman E, Peterka T, and Altintas I et al. The future of scientific workflows. The International Journal of High Performance Computing Applications, 32(1):159– 175, 2018.

[12] Liu W, Jia J, and Dai Y, et al.. Delineating COVID-19 immunological features using single-cell RNA sequencing. The Innovation, 3(5):100289, 2022.

[13] Aznaourova M., Schmerer N, and Janga H et al. Single-cell RNA sequencing uncovers the nuclear decoy lincRNA PIRAT as a regulator of systemic monocyte immunity during COVID-19. PNAS, 119(36):1–12, 2022.

[14] Schulte-Schrepping J, Reusch N, and Paclik D et al. Severe covid-19 is marked by a dysregulated myeloid cell compartment. Cell, 182(6):1419–1440, 2020.

[15] Yao C, Bora SA, and Parimon T et al. Cell-type-specific immune dysregulation in severely ill covid-19 patients. Cell Reports, 34(1), 2020.

[16] Song E, Bartley CM, and Chow RD. Divergent and self-reactive immune responses in the cns of covid-19 patients with neurological symptoms. Cell Reports Medicine, 2(5), 2021.

[17] Silvin A, Chapuis N, and Dunsmore G et al. Elevated calprotectin and abnormal myeloid cell subsets discriminate severe from mild covid-19. Cell, 182(6), 2020.

[18] Guo C, Li B, and Ma H et al. Tocilizumab treatment in severe COVID-19 patients attenuates the inflammatory storm incited by monocyte centric immune interactions revealed by single-cell analysis. Nature Communications, 11:3924, 2020.

[19] Jørgensen MJ, Holter JC, and Christensen EE et al. Increased interleukin-6 and macrophage chemoattractant protein-1 are associated with respiratory failure in covid-19. Scientific reports, 10(1):1–11, 2020.

[20] Stephenson E, Reynolds G, and Botting RA et al. Single-cell multi-omics analysis of the immune response in COVID-19. Nature Medicine, 27:904–916, 2021.

[21] Zhang X, Li T, and Liu F et al. Comparative analysis of droplet-based ultrahigh-throughput single-cell rna-seq systems. Molecular cell, 73(1):130–142, 2019.

[22] Tom Smith, Andreas Heger, and Ian Sudbery. Umi-tools: Modeling sequencing errors in unique molecular identifiers to improve quantification accuracy. Genome Research, 27:gr.209601.116, 01 2017.

[23] S Parekh, C Ziegenhain, B Vieth, W Enard, and I Hellmann. zUMIs A fast and flexible pipeline to process RNA sequencing data with UMIs. GigaScience, 7(6), 05 2018.

[24] Hao Y, Hao S, and Andersen-Nissen E et al. Integrated analysis of multimodal single-cell data. Cell, 2021.

[25] Young MD and Behjati S. Soupx removes ambient rna contamination from droplet-based single-cell rna sequencing data. Gigascience, 9(12):giaa151, 2020.

[26] Luecken MD and Theis FJ. Current best practices in single-cell RNA-seq analysis: a tutorial. Molecular Systems Biology, 15(e8746):1–23, 2019.

[27] Heimberg G, Bhatnagar R, and El-Samad H et al. Dimensionality in Gene Expression Data Enables the Accurate Extraction of Transcriptional Programs from Shallow Sequencing. Cell Systems, 2(4):239–250, 2016.

[28] Stuart T, Butler A, and Hoffman P et al. Comprehensive integration of single-cell data. Cell, 177(7):1888–1902, 2019.

[29] Tran HTN, Ang KS, and Chevrier M et al. A benchmark of batch-effect correction methods for single-cell rna sequencing data. Genome biology, 21(1):1–32, 2020.

[30] Becht E, McInnes L, and Healy J et al. Dimensionality reduction for visualizing single-cell data using umap. Nature biotechnology, 37(1):38–44, 2019.

[31] Ringńer M. What is principal component analysis? Nature biotechnology, 26(3):303–304, 2008.

[32] Baran Y, Bercovich A, and Sebe-Pedros A et al. Metacell: analysis of single-cell rna-seq data using k-nn graph partitions. Genome biology, 20(1):1–19, 2019.

[33] Hao Y, Hao S, and Andersen-Nissen E et al. Integrated analysis of multimodal single-cell data. Cell, 2021.

[34] Zhang AW, O’Flanagan C, and Chavez EA et al. Probabilistic cell-type assignment of single-cell rna-seq for tumor microenvironment profiling. Nature methods, 16(10):1007–1015, 2019.

[35] Kingma DP and Ba J. Adam: A method for stochastic optimization. *arXiv preprint arXiv:1412*.6980, 2014.

[36] Abadi M, Barham P, and Chen J aet al.. *{*TensorFlow*}*: a system for *{*LargeScale*}* machine learning. In 12th USENIX symposium on operating systems design and implementation (OSDI 16), pages 265–283, 2016.

[37] Ianevski A, Giri AK, and Aittokallio T. Fully-automated and ultra-fast cell-type identification using specific marker combinations from single-cell transcriptomic data. Nature communications, 13(1):1–10, 2022.

[38] Aran D, Looney AP, and Liu L et al. Reference-based analysis of lung single-cell sequencing reveals a transitional profibrotic macrophage. Nature immunology, 20(2):163–172, 2019.

[39] Shao X, Liao J, and Lu X et al.. sccatch: automatic annotation on cell types of clusters from single-cell rna sequencing data. Iscience, 23(3):100882, 2020.

[40] Herring CA, Banerjee A, and McKinley ET et al. Unsupervised Trajectory Analysis of Single-Cell RNA-Seq and Imaging Data Reveals Alternative Tuft Cell Origins in the Gut. Cell Systems, 6(1):37–51, 2018.

[41] Street K, Risso D, and Fletcher R et al. Slingshot: cell lineage and pseudotime inference for single-cell transcriptomics. BMC Genomics, 19(477):1–16, 2018.

[42] Wolf FA, Hamey FK, and Plass M et al. PAGA: graph abstraction reconciles clustering with trajectory inference through a topology preserving map of single cells. Genome Biology, 20(59):1–9, 2019.

[43] Dimitrov D, Türei D, and Garrido-Rodriguez M, et al.. Comparison of methods and resources for cell-cell communication inference from single-cell rna-seq data. Nature Communications, 13(1):1–13, 2022.

[44] Efremova M, Vento-Tormo M, and Teichmann SA et al. Cellphonedb: inferring cell–cell communication from combined expression of multi-subunit ligand– receptor complexes. Nature protocols, 15(4):1484–1506, 2020.

[45] Shao X, Liao J, and Li C et al. Celltalkdb: a manually curated database of ligand–receptor interactions in humans and mice. Briefings in bioinformatics, 22(4):bbaa269, 2021.

[46] Nöel F, Massenet-Regad L, and Carmi-Levy I, et al.. Dissection of intercellular communication using the transcriptome-based framework icellnet. Nature communications, 12(1):1–16, 2021.

[47] Cillo AR, Kürten CHL, and Tabib T, et al.. Immune landscape of viral-and carcinogen-driven head and neck cancer. Immunity, 52(1):183–199, 2020.

[48] Jin S, Guerrero-Juarez CF, and Zhang L et al. Inference and analysis of cell-cell communication using cellchat. Nature communications, 12(1):1–20, 2021.

[49] Palla G, Spitzer H, and Klein M et al. Squidpy: a scalable framework for spatial omics analysis. Nature methods, 19(2):171–178, 2022.

[50] Liberzon A, Subramanian A, and Pinchback R et al. Molecular signatures database (MSigDB) 3.0. Bioinformatics, 27(12):1739–1740, 2011.

[51] Fabregat A, Jupe S, and Matthews L et al. The Reactome Pathway Knowledgebase. Nucleic Acids Research, 4(46(D1)):D649–D655, 2018.

[52] Mi H, Ebert D, and Muruganujan A et al. PANTHER version 16: a revised family classification, tree-based classification tool, enhancer regions and extensive API. Nucleic Acids Research, 49(D1):D394–D403, 2020.

[53] Huang D, Sherman B, and Lempicki R. Systematic and integrative analysis of large gene lists using DAVID bioinformatics resources. Nature Protocols, 4:44–57, 2009.

[54] Griss J, Viteri G, and Sidiropoulos K et al. ReactomeGSA Efficient Multi-Omics Comparative Pathway Analysis. Molecular and Cellular Proteomics, 19(12):2115– 2125, 2020.

[55] Damiani C, Maspero D, and Di Filippo M et al. Integration of single-cell rna-seq data into population models to characterize cancer metabolism. PLoS computational biology, 15(2):e1006733, 2019.

[56] Wagner A, Wang C, and Fessler J et al. Metabolic modeling of single th17 cells reveals regulators of autoimmunity. Cell, 184(16):4168–4185, 2021.

[57] Alghamdi N, Chang W, and Dang P et al. A graph neural network model to estimate cell-wise metabolic flux using single-cell rna-seq data. Genome research, 31(10):1867–1884, 2021.

[58] Wu Y, Yang S, and Ma J et al. Spatiotemporal immune landscape of colorectal cancer liver metastasis at single-cell levelspatial and cellular landscape of crlm. Cancer discovery, 12(1):134–153, 2022.

[59] Clough E and Barrett T. The gene expression omnibus database. Methods in molecular biology, 1418:93–110, 2016.

[60] R Core Team. *R: A Language and Environment for Statistical Computing*. R Foundation for Statistical Computing, Vienna, Austria, 2021.

[61] Däeron M. Fc receptors as adaptive immunoreceptors. Fc Receptors, pages 131– 164, 2014.

[62] van de Veerdonk FL, Giamarellos-Bourboulis E, and Pickkers P et al. A guide to immunotherapy for covid-19. Nature Medicine, 28(1):39–50, 2022.

[63] Bournazos S, Gupta A, and Ravetch JV. The role of igg fc receptors in antibody-dependent enhancement. Nature Reviews Immunology, 20(10):633–643, 2020.

[64] Däeron M, Latour S, and Malbec O, et al.. The same tyrosine-based inhibition motif, in the intra-cytoplasmic domain of fc*γ*riib, regulates negatively bcr-, tcr-, and fcr-dependent cell activation. Immunity, 3(5):635–646, 1995.

[65] Ben Mkaddem S, Benhamou M, and Monteiro RC. Understanding fc receptor involvement in inflammatory diseases: from mechanisms to new therapeutic tools. Frontiers in immunology, 10:811, 2019.

[66] Foss S, Bottermann Ma, and Jonsson Al et al. Trim21—from intracellular immunity to therapy. Frontiers in Immunology, 10:2049, 2019.

[67] Pyzik M, Sand KMK, and Hubbard JJ et al. The neonatal fc receptor (fcrn): a misnomer? Frontiers in immunology, page 1540, 2019.

[68] Suzuki T, Kono H, and Hirose N et al. Differential involvement of src family kinases in fc*γ* receptor-mediated phagocytosis. The Journal of Immunology, 165(1):473–482, 2000.

[69] Däeron M. Fc receptor biology. Annual review of immunology, 15:203, 1997.

[70] Mócsai Aand Ruland J and Tybulewicz VLJ. The syk tyrosine kinase: a crucial player in diverse biological functions. Nature Reviews Immunology, 10(6):387–402, 2010.

[71] Klippel A, Reinhard C, and Kavanaugh WM et al. Membrane localization of phosphatidylinositol 3-kinase is sufficient to activate multiple signal-transducing kinase pathways. Molecular and cellular biology, 16(8):4117–4127, 1996.

[72] Gilfillan AM and Tkaczyk C. Integrated signalling pathways for mast-cell activation. Nature Reviews Immunology, 6(3):218–230, 2006.

[73] Koretzky GA, Abtahian F, and Silverman MA. Slp76 and slp65: complex regulation of signalling in lymphocytes and beyond. Nature Reviews Immunology, 6(1):67–78, 2006.

[74] Neumann K and Ruland J. Kinases conquer the inflammasomes. Nature immunology, 14(12):1207–1208, 2013.

[75] Kong X, Liao Y, and Zhou L et al. Hematopoietic cell kinase (hck) is essential for nlrp3 inflammasome activation and lipopolysaccharide-induced inflammatory response in vivo. Frontiers in pharmacology, 11:581011, 2020.

[76] Petrilli V, Papin S, and Tschopp J. The inflammasome. Current Biology, 15(15):R581, 2005.

[77] Liu T, Zhang L, and Joo D et al. Nf-*κ*b signaling in inflammation. Signal transduction and targeted therapy, 2(1):1–9, 2017.

[78] Yan Z, Luo H, and Xie B et al. Targeting adaptor protein slp76 of rage as a therapeutic approach for lethal sepsis. Nature communications, 12(1):1–14, 2021.

